# Investigation of the effect of estradiol and subculturing on the receptor expression within hormone-dependent breast cancer cells

**DOI:** 10.1101/548271

**Authors:** Avijit Mallick, Sabbir Ahmed

**Affiliations:** School of Science and Sport, University of the West of Scotland; Department of Pharmacy and Biomedical Sciences, University of Portsmouth

**Author notes:** Current address: Department of Biology, McMaster University (Avijit Mallick). Corresponding Author: Avijit Mallick, 1280 Main St. West, Hamilton, ON L8S4K1, Canada.

**Keywords:** breast cancer, estradiol, estrogen receptor, apoptosis

## Abstract

It is now known that a very crucial role in breast cancer development, prognosis and occurrence is played by the estrogen receptor (ER). The steroid hormone estradiol (E2) acts via two nuclear receptors, estrogen receptor-α (ERα) and estrogen receptor-β (ERβ). E2 was shown previously to increase breast cancer cell proliferation in a dose-dependent manner and also induce apoptosis in long term estrogen deprived breast cancer cells. Studies have also shown that the degree of subculturing affects cell line property including gene expression. The aim of this study was to investigate the effect of E2 concentration on cell proliferation, morphology and ER expression and to investigate the effect of subculturing on the expression of ER. Our results have shown that an increase in E2 concentration was found to increase MCF-7 cell proliferation, but extreme concentrations caused significantly low cell proliferation and induced apoptosis. Moreover, ERα expression was significantly upregulated with an increase in E2 concentration, whereas ERβ2 expression was found to be unchanged at low E2 concentration and significantly upregulated at higher E2 concentration. ERα expression at passage 3 ([E2]=1nM) was significantly downregulated compared to the cells at passage 0, in addition to the significant downregulation of the same at E2 concentrations of 1nM and 10µM compared to the untreated control sample. Overall, our data suggests that high concentration of E2 can reduce proliferation and induce apoptosis in the breast cancer cells. Increased E2 exposure and subculturing also appear to change the ERα expression significantly in the breast cancer cell line.

## INTRODUCTION

### Breast cancer

Breast cancer is the leading type of cancer in women accounting for approximately 25% of all cases of cancer resulting in 1.68 million cases and 522,000 deaths worldwide in 2012 (Stewart and Wild, 2014). Breast cancer is the most frequently diagnosed cancer among women and was first shown to be estrogen-dependent by Beatson (1896) who subsequently showed the beneficial effects of oophorectomy in pre-menopausal patients.

Invasive breast cancers are often classified based on the presence or absence of hormone receptors and the presence or lack of HER2, an epidermal growth factor receptor family protein. Hormone positive breast cancer cells contain either estrogen receptors (ER) or progesterone receptors (PR) or both. Approximately 67% of breast cancers have been shown to contain either ER or PR, the percentage has been shown to be higher for older women (Lund *et al.,* 2010). Hormone therapy drugs that either lower estrogen levels or block ER have been shown to be effective against estrogen-dependent breast cancer but are not helpful for hormone receptor negative breast cancer cells which have been shown to grow faster than the hormone receptor-positive cells. On the other hand, breast cancers with high epidermal growth factor receptor family protein HER2/neu expression, which account for about 20% of all breast cancer cases, are termed as HER2 positive whereas those without high expression of HER2 as HER2 negative. Lastly, triple positive breast cancers with ER, PR and high expression of HER2, grow and spread faster than most of the other types and are more frequent in younger women (Lund *et al*., 2010).

### Estrogen receptors

The estrogen family of steroids play vital roles in the development and maintenance of sexual and reproductive function. Moreover, they exert various biological effects on the central nervous, musculoskeletal, cardiovascular and immune system in both sexes (Gustafsson, 2003). 17β-Estradiol (E2), the most potent estrogen, acts by binding to two ER subtypes: ERα and ERβ which possess distinct and essential roles. The two types of ERs are encoded by the genes Estrogen receptor 1 (ESR1) and Estrogen receptor 2 (ESR2) respectively on the fifth and fourteenth human chromosomes. Hormone activated ER form dimers with each other: ERα (αα), ERβ (ββ) homodimers or ERαβ (αβ) heterodimers (Tremblay *et al*., 1997). Although these ERs bind to identical DNA response elements, different effects, by transcriptional regulation of the target genes, are exerted on the morphology, motility and proliferation of cells (Lazennec *et al.* 2001).

### ER subtypes

Of the proteins expressed, ERα is the main protein expressed by the breast cancer cells, a cytoplasmic/nuclear receptor that activates upon ligand binding or phosphorylation and regulates the transcription of its targeted genes by binding to the estrogen response element (ERE) or interacting with other transcription factors upstream of the targeted genes (Heldring *et al*., 2007). ERα expression and breast cancer biology is shown to have a high level of correlation, with the breast carcinomas lacking ERα expression often showing more aggressive phenotypes. As a pathological diagnostic criterion in breast cancer, immunohistochemical staining has shown that positive staining of ERα protein expression in the nucleus act as a good prognostic indication. It is also being found to be localized in the cytoplasm and/or membrane.

ERβ was identified in 1996 (Kuiper *et al.,* 1996) and has also been shown to be expressed in various cancer cells including breast cancer cells (Omoto *et al*., 2002). Studies have shown that ERβ can antagonize ERα dependent transcription, with recruitment of c-Fos to AP-1 regulated promoters (Matthews and Gustafsson 2003; Matthews *et al*., 2006). Furthermore, ERα proteolytic degradation is increased by the expression of ERβ and its variant ERβ2 suggesting that ERβ mediated inhibition of cell proliferation is accomplished by the combination of key transcription factor recruitment and ERα degradation (Matthews *et al*., 2006). ERβ is found to be expressed in ERα positive breast cancers and immunohistochemical studies have also shown the presence of both these receptors in human breast cancer cells (Leygue *et al*., 1999). ERβ expression is usually higher than that of ERα in the breast cancer cells, but much lower when compared to healthy tissue. ERβ subtypes can only transactivate transcription when heterodimer has been formed with ERβ1, but ERβ2 (with significantly reduced E2 binding and ERE binding ability) preferentially forms a dimer with ERα thus inhibiting its DNA binding (Iwase *et al*. 2003). Studies suggest that a balance between ERα and ERβ signaling dictates the overall response of proliferation induced by E2. ERβ1 and ERβ2/ERβcx specifically suppress the function of ERα, particularly proliferation, through various pathways and thus a change in the ERα: ERβ ratio may lead to the progression and development of tumours (Leygue *et al.* 1999).

### Cell proliferation and subculturing

Cell division in the body is highly regulated by several signaling pathways, but when these signals are faulty or missing, cells start to proliferate leading to tumour formation. Underlying this outcome is the accumulated change in genes controlling the cell division by the means of gene duplication or alterations by substitution, deletion or addition of nucleotide bases in DNA leading to uncontrolled cell proliferation. To combat this, cells have repair mechanisms, but the balance between mutations and repair gradually shifts with time allowing the mutations to accumulate prior to the cells becoming cancerous.

Similarly, growing evidence from studies have shown that the passage number, degree of subculturing or the number of times the cells being transferred into new vessel with confluence of ≈80%, affects it characteristics with time (Esquenet *et al*., 1997; Briske-Anderson *et al*. 1997; Chang-Liu *et al.* 1997). Likewise, cancer cell lines experience changes in response to stimuli, protein expression, transfection efficiency, morphology and growth rates at higher passage number. Also, cells in a culture are under environmental and manipulative stress leading to the subject of evolutionary process of competition and natural selection. As such, most cell cultures represent heterogeneous mix of cells competing for resources such as growth factors, nucleic acids and salts. These cells when given an advantage (e.g faster growth factor) outcompete other types of cells giving rise to a different population set. Culturing of primary tumour cells is sometimes difficult, so transformed cell lines with well characterized cytogenetics and biochemical markers available commercially are used in the laboratories. The transformed and diseased cell lines are of special concern because in the abnormal starting population evolutionary changes take place at a faster rate in both the genotypic and phenotypic levels. Moreover, one or all of the cellular check point genes (such as p53, pRB) in these cell lines are altered in parallel to other mutations and are thus prone to increased genetic instability with continuous subculturing.

### Basis of present investigation

A study analyzing the effect of the four estrogens: estrone (E1), E2, estriol (E3) and estetrol (E4) on the proliferation and ERα/ERβ expression of breast cancer cell line ZR 75-1 showed that E1 and E4 have low stimulation of cell proliferation at lower concentration; all the estrogens (E2, E3 and E4) except E1 had significantly lower proliferative action at high concentration. Although ERα expression was significantly increased by all the estrogens (with significant difference between each of them), none of the estrogens could significantly change the ERβ expression compared to the control (Li *et al*. 2015). It has been shown previously that E2 promotes MCF-7 cell proliferation in a dose-dependent manner where the highest concentration used was 100nM (Pattarozzi *et al*., 2008). In contrast, a study by Altiok *et al*. (2006) showed that E2 induced apoptosis (IC_50_=2µM) in dose-dependent manner under low growth condition. E2 was not however, cytotoxic in the range of E2 concentration used at higher level of FBS, though the data supporting this was not presented (Altiok *et al*., 2006). However, several studies have shown that even low concentration of E2 can induce apoptosis in long term estrogen deprived breast cancer cells (Jordan *et al*., 2005; Song *et al*., 2001). It has also been shown that E2 enhanced proliferation with increasing concentration from 1nM to 1µM, however, at higher concentration (10µM) proliferation was found to decrease (Ma *et al*., 2013). But no studies have yet analyzed the effect of varied E2 doses and the passage number of the cell lines on their receptor expression. In this study, we consider the impact on cell proliferation and receptor expression of high E2 exposure. Additionally, the effect of subculturing was also be investigated with high concentration of E2 so as to investigate the effect of E2 on the targeted receptor expression ERα and ERβ at different passage number.

## MATERIALS AND METHODS

### Preparation of media

Media was prepared by adding 50mL FBS, 5mL penicillin/streptomycin mix and 5mL Fungizone Amphotericin into each of the 500mL phenol red free and phenol red DMEM containing 4.5g/L D-glucose, L-glutamine and pyruvate.

### Preparation of E2 stock

E2 stock of 10mM was prepared in 1mL DMSO which was then serially diluted to give stocks of 0.1mM and 1µM and stored at −20°C.

### Preparation of reagents and cell staining

Paraformaldehyde, 4% w/v, was prepared by dissolving 4g of paraformaldehyde in 100mL of DPBS at 60°C. Sodium hydroxide solution of 1M concentration was then prepared by dissolving 0.4g in 10mL which was then added dropwise into the paraformaldehyde solution until the pH was 7.4. Triton X-100, 0.2% v/v, was prepared by adding 100µL of Triton X-100 into 50mL double distilled water. DAPI stock was prepared by dissolving 1mg of DAPI in 1mL DMSO. It was then further diluted by 100 fold in glycerol.

Cells were rinsed three times with 1mL DPBS. After incubating in 1mL 4% paraformaldehyde for 10min cells were then rinsed again in 1mL DPBS for three times each lasting 5min. Following this, cells were permeabilized by immersing in 1mL 0.2% Triton X-100 for 5min and rinsed twice with 1mL DPBS. Three glass slides were prepared with a drop of DAPI solution for each sample and the cover slip was then placed over the drop upside down using forceps. The slides were kept in the dark overnight and viewed under the fluorescence microscope Olympus IX71 at a magnification of X20 and images taken using Olympus DP70.

### Cell Cultures

MCF-7 cell line was maintained in T-75 flasks with phenol red DMEM containing 10% heat inactivated FBS, 1% Fungizone and 1% penicillin/streptomycin at 37°C in a humidified 5% CO_2_ atmosphere. For subculturing, on reaching a confluence of ≈80%, the media was removed from the flask and 2mL DPBS was added. After discarding the DPBS, 2mL trypsin was added and the flask was kept in the incubator for 2min at 37°C in a humidified 5% CO_2_ atmosphere after which 3mL phenol red free DMEM was added and mixed thoroughly. The cell suspension was then placed in a fresh tube and centrifuged at 1200rpm for 5min. The pellet was then re-suspended in 1mL of phenol red free DMEM and seeded into two new T-75 flask containing 15mL phenol red DMEM, each with splitting ratio of 1:2 by adding 500µL of the cell suspension. The flasks were then incubated at 37°C in a humidified 5% CO_2_ atmosphere.

### Cell counting

On reaching a confluence of ≈80%, cells from the flask were placed in a fresh tube and 1mL of cell suspension made following the procedure described above. The cell suspension of 10µL was added to 20µL trypan blue solution in a separate tube and mixed thoroughly, placed under the glass slide on the haemocytometer and cells counted under light microscope to determine the cell concentration.

### Cell proliferation study

Six well plates were set up with MCF-7 cell concentrations of 1X10^5^ cells/mL/well in 4mL phenol red free DMEM with E2 concentrations of 1nM, 10nM, 100nM, 1µM, 10µM, 100µM and control in duplicate. Cells were allowed to adhere for 24h before being exposed to E2. After 4 days of incubation at 37°C in a humidified 5% CO_2_ atmosphere, media from the wells was removed and 600µL trypsin added for 2min followed by 1mL phenol red free DMEM and mixed thoroughly. The cell suspension was placed in a new tube and centrifuged at 1200rpm for 5min. The pellet was then re-suspended in 1mL media from which 10µL of the cell suspension was added to 20µL trypan blue solution in a separate tube and cell count determined following the procedure described above.

### Sub culturing

Six well plates with MCF-7 cells at concentration of 10^5^cells/mL/well were set up in duplicate with E2 concentrations of 1nM, 10µM and untreated control sample. The plates were kept in the incubator at 37°C in a humidified 5% CO_2_ atmosphere and sub cultured every 4days to a new six well plate with an initial seeding concentration of 1X10^5^cells/mL/well.

### RNA extraction

From the 1mL cell suspension in the 5mL tube 5X10^5^ cells were transferred into a new 1.5mL tube and 1mL TRIzol® Reagent was added after which the cells were lysed and total RNA extracted following the protocol provided by the manufacturer. Consequently, RNA quantity and quality was assessed using a NanoDrop ND-1000 UV-Vis Spectrophotometer.

### cDNA synthesis and qRT-PCR

Using the QuantiTect Reverse Transcription Kit total RNA was converted to cDNA after the elimination of any genomic DNA following the manufacture’s instruction. cDNA sample was diluted (50%) by adding 20µL of RNase free water and mixed thoroughly.

Real-time PCR with SYBR Green fluorescent marker and specific primers (Table 1) were carried out. Frozen cDNA samples were thawed, vortexed and centrifuged before setting up duplication of each reaction in 96-well reaction plate with 0.2µL of each primer (0.1µL each of forward and reverse), 2µL of cDNA, 6µl of Fast SYBR® Green Master Mix and 3.8µL of RNase free water to make up a total reaction volume of 12µL. The plate was then covered tightly with a cover slip followed by PCR in StepOnePlus™ Real-Time PCR System with the incubation times as follows: 95°C for 10min followed by 40 cycles of 95°C for 15s and 60°C for 1min.

**Table 1:**
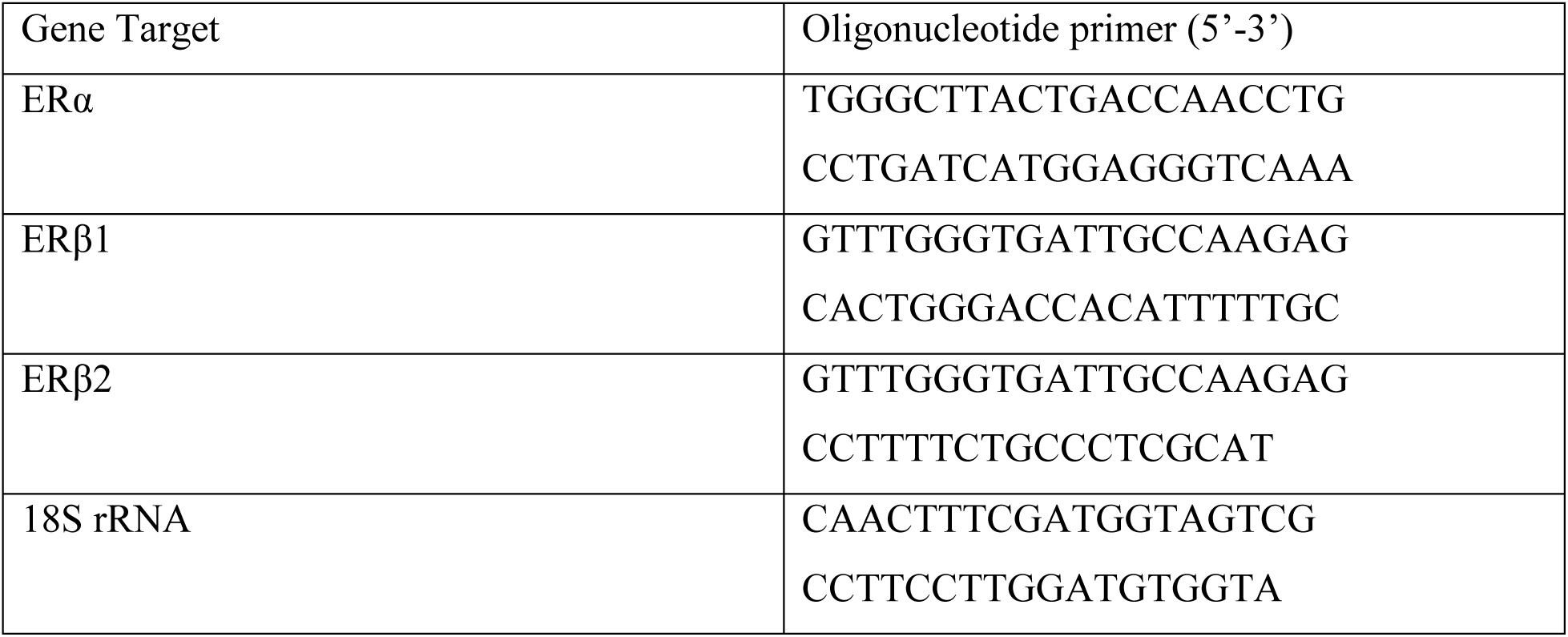
List of primers used.

### Statistical analysis

The quantitative data were presented as mean ± standard error of mean (SEM). Statistical analysis was performed by means of analysis of variance (ANOVA) followed by Student t-test with differences considered significant at a *p*-value of <0.05.

## RESULTS AND DISCUSSION

### High E2 concentration increases cell proliferation only during initial incubation

To evaluate the effect of time dependent E2 exposure, cells were incubated for 4, 7 and 10 days at an E2 concentration of 1µM (cells treated at this [E2] gave highest proliferation in a study by Ma et al. (2013)). It was found that the cell count at the end of 4 days incubation was significantly higher (\**p* <0.05 vs. unstimulated control sample; Fig. 1) than the control. The rate of proliferation was found to continue beyond the 7^th^ day resulting in both the control and E2 treated wells containing almost equal number of cells after 10 days (Fig. 1). The results also showed that the cell proliferation increased in a time dependent manner up to the 7^th^ day. Though the cells in the E2 treated wells was equal to that of the control on the 10^th^ day, the space available in the wells was found to be a limiting factor as the cells in both the wells had 100% confluence. Incubation time of 4 days was therefore considered as the optimum time for the proliferation to have taken place so as to provide enough cells for the RNA extraction as well as allowing further sub culturing of cells to be undertaken.

**Figure 1:**
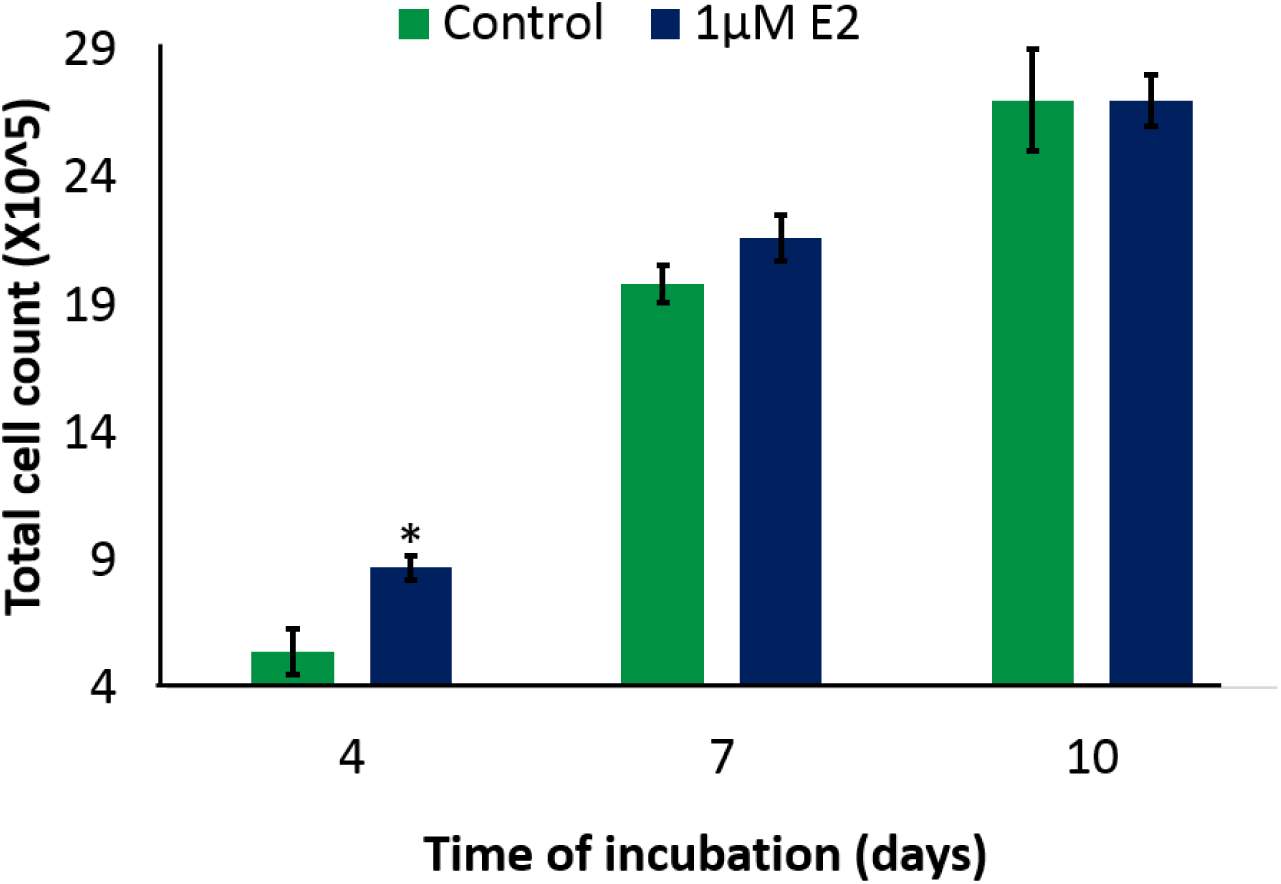
Effect of incubation time on cell proliferation. Cell proliferation at 1µM E2 concentration is significantly higher after 4 days incubation, but then show similar proliferation after 7 and 10 days of incubation. The error bars indicate standard error of mean, and two-tailed Student’s t-tests were used to determine the statistical significance: \**p* < 0.05 (n=3).

### MCF-7 cell line with higher passage number shows increased cell proliferation

The effect of increasing concentration of E2 on ER positive MCF-7 cell line of different passage number was then evaluated using trypan blue exclusion assay where the transparent cells under the microscope were counted as viable. MCF-7 cell lines of different passage numbers were used, one of which was recently purchased at passage number of around 150 and the other with passage number of around 500 was in use in the lab for at least one year.

It was observed that an increase in E2 concentration from 1nM to 10nM resulted in an increased cell proliferation in a concentration-dependent manner at 4^th^ day of incubation by approximately 55% and 60% in the old (passage number of ∼500) and new cell line (passage ∼150) compared to the untreated control (Fig. 2). Interestingly, further increase in E2 concentration decreased cell proliferation by approximately 88% and 74% in both the old and new cell lines at 100µM E2 concentration. Moreover, the cell proliferation rate in the old cell line was higher than the new cell line in each of the samples except at 100µM.

**Figure 2:**
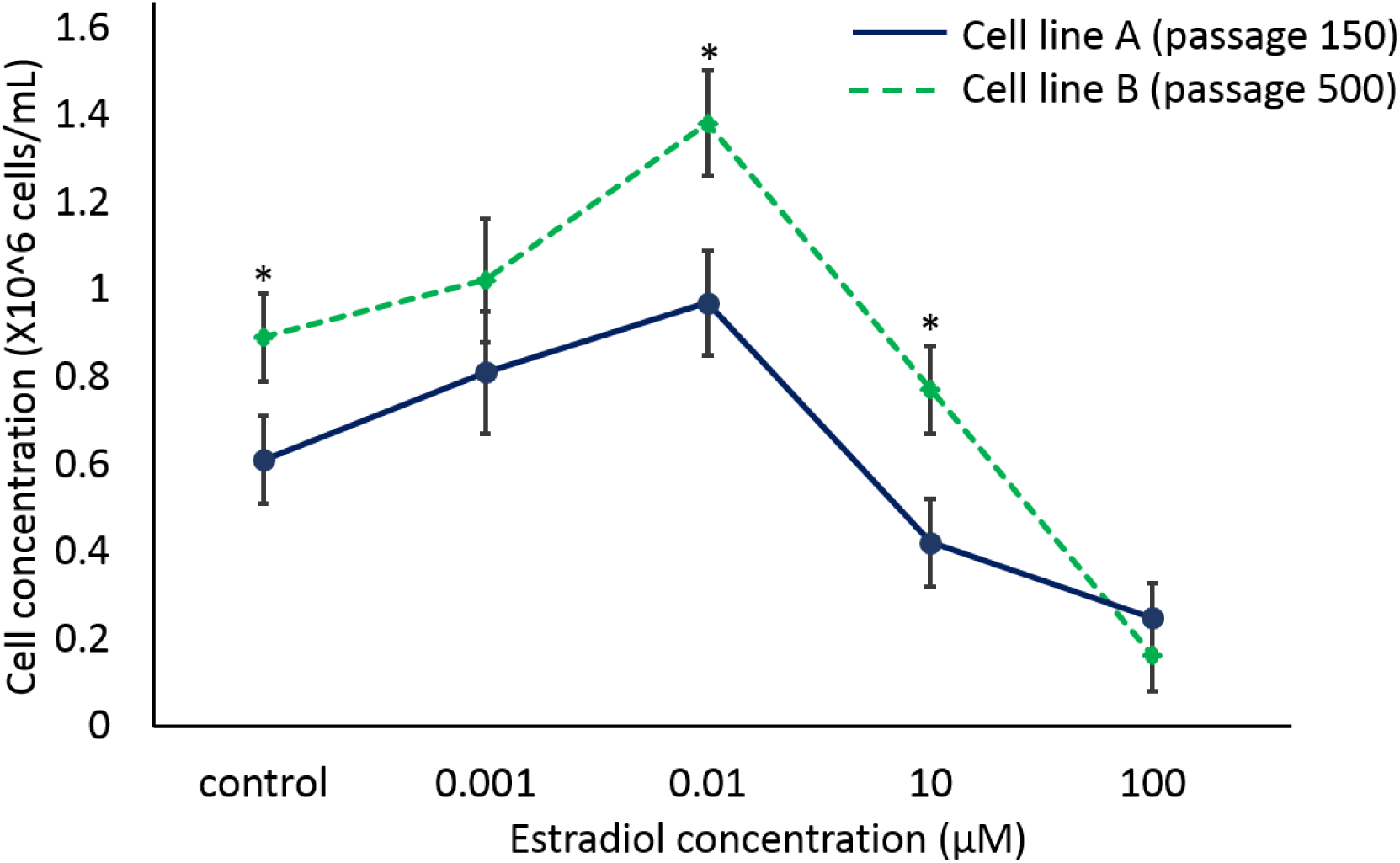
Effect of E2 on two different MCF-7 cell line proliferation with varied passage numbers. Data showing increasing E2 concentration effect on two different MCF-7 cell line proliferation after 4 days. The error bars indicate standard deviation, and two-tailed Student’s t-tests were used to determine the statistical significance: \*\**p* < 0.01 (n=2).

As previously discussed, cells accumulate mutation with time and proliferate at a high rate, a main characteristic of cancer cells. Concordant to the theory, the old cell line could be assumed to have had a considerable higher proliferation rate compared to the new cell line with both the cell lines having the highest rate of cell proliferation at E2 concentration of 0.01µM. Due to the change in protocol, we then repeated the experiment with the new cell line and two more E2 concentrations of 0.1µM and 1µM compared to the above experiment. Cells were therefore incubated in duplicate for 24h before being exposed to E2 thus allowing them to adhere to the surface of the wells. Similar to the previous finding, it was found that the increase in E2 concentration increased cell proliferation in a dose-dependent manner, but this time the peak of cell growth was at 1µM (Fig. 2). E2 bound to ERα increases the gene expression required for cell proliferation and with the increase in E2 concentration more ERαs are present in the complex form (E2-ERα) thus increasing the downstream signaling. Similar to Figure 2, further increase in E2 concentration decreased cell proliferation and inhibited cell growth. Cell growth at 1µM was significantly increased and at 100µM was significantly reduced compared to the untreated control sample (\**p* < 0.05 vs. unstimulated control sample; Fig. 3). In addition, cell growth at 100µM was also significant reduced compared to the 1µM treated sample (^+^*p* < 0.05 vs. 100µM sample; Figure 3).

**Figure 3:**
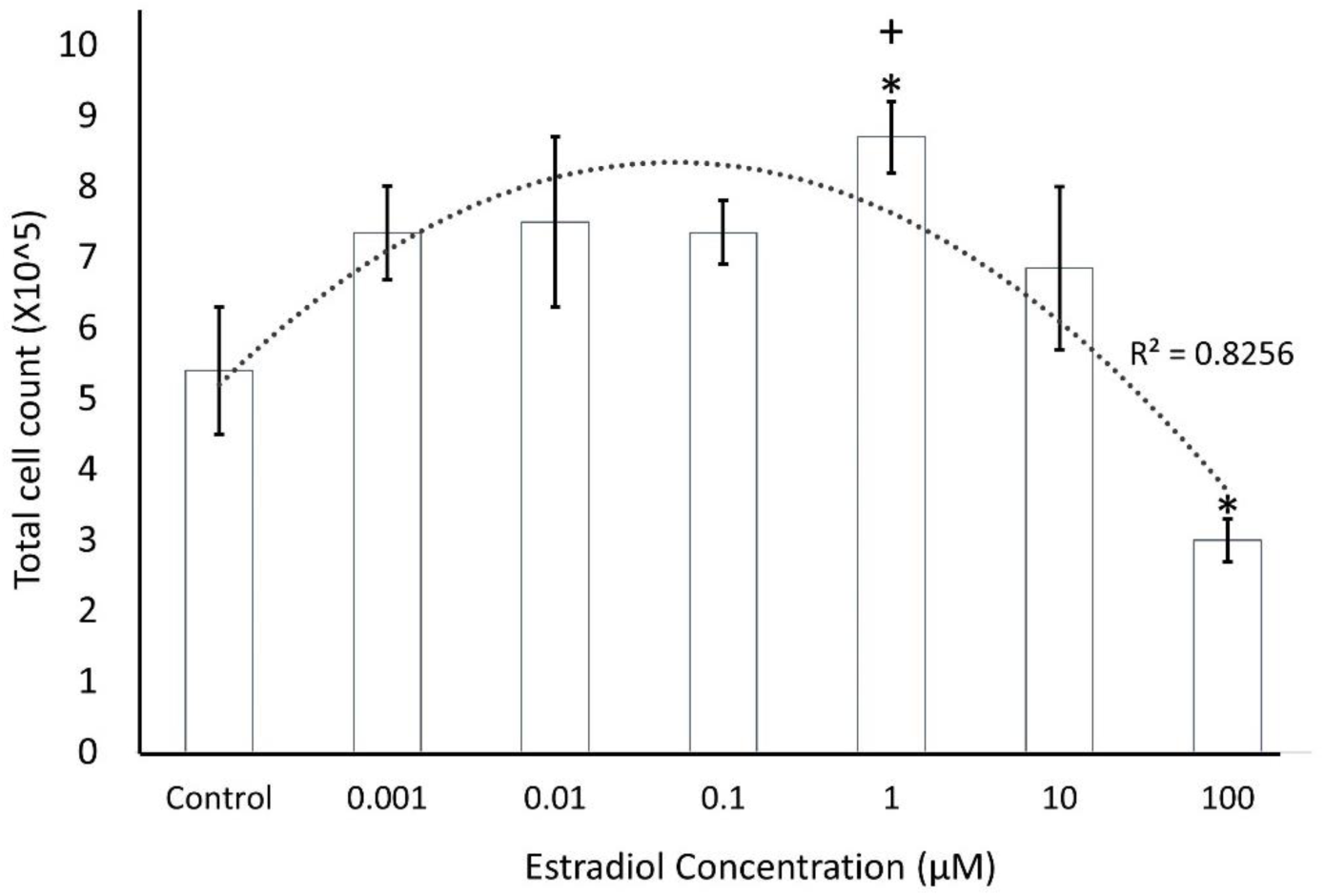
High E2 concentration is detrimental for the cancerous cells. Data showing increased proliferation of MCF-7 cells with increase in concentration of E2 and then decrease in the total cell number at high concentration of 100µM. The error bars indicate standard error of mean, and two-tailed Student’s t-tests were used to determine the statistical significance: (\**p* < 0.05 vs. unstimulated control sample; ^+^*p* < 0.05 vs. 100µM sample, n=2).

E2 promotes mammary gland cell proliferation by changing hormone responsive gene expression involved in the cell cycle and/or cell death. It has been discussed previously how ERα receptor signaling increases cell proliferation by regulating gene expression. More importantly, E2 has also been shown as a potent apoptotic inhibitor in MCF-7, ZR-75-1 and T47-D breast cancer cells where it regulates the apoptotic gene expression including Bcl-2 (Gompel *et al.,* 2000). Moreover, the above data shows that extremely high concentration of E2 significantly inhibits the cell proliferation contradicting its proliferative role.

### Apoptosis at high E2 concentration

It has been shown above that at high concentration E2 inhibited cell proliferation and to further evaluate any morphological change, cells were viewed under light microscope before and after E2 exposure and images taken (Fig. 4A). There was no change observed in the morphology of the control sample cells but there was clear difference between the cells in the 100µM sample before and after 24h of E2 exposure. The image after 24h of incubation at 100µM indicated apoptosis as circular bodies were clearly seen. As a main characteristic of apoptosis, cells break up into apoptotic bodies containing fragmented DNA and organelles (Kerr *et al*., 1972). To support the occurrence of apoptosis, cells were stained with DAPI and visualized under fluorescent microscope. Untreated cells and cells exposed to E2 (10nM and 1µM) were found to possess normal morphology with intact nuclei whereas the cells at higher E2 concentrations (10µM and 100µM) exhibited the characteristics of apoptosis with condensed and fragmented nuclei (Figure 4B, C). The proportion of apoptotic cells were found to be higher at 100µM than 10µM thus supporting the above cell proliferation data.

**Figure 4:**
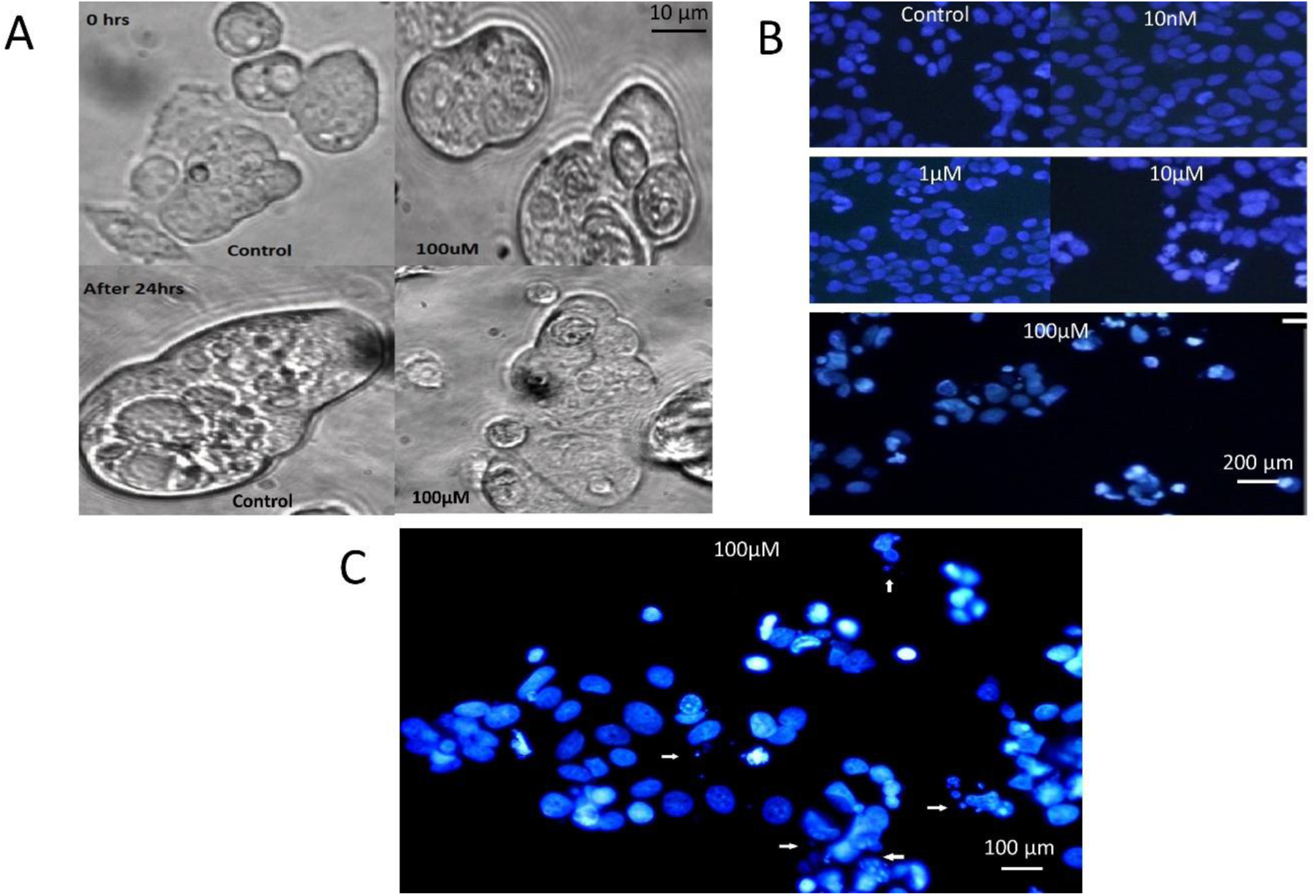
Cells undergo apoptosis at E2 concentration of 100µM. **A)** Cells under light microscope(X40) before and after E2 exposure. Showing the cell progression status at 0 and 24hrs for both untreated and treated cells with 100µM E2. **B)** Morphological assessment of MCF-7 cells with DAPI staining at four different E2 concentrations. Representative images for control cells, cells exposed to 10nM, 1µM, 10µM and 100µM E2 concentrations. **C)** Showing apoptosis in MCF-7 cells treated with 100µM E2.

Another property of E2 action exist, i.e. inducing apoptosis, which contrasts its ability to promote cell proliferation and therefore inhibit apoptosis, revealed once again by the observed data. It has been shown previously by several studies that E2 is also capable of inducing apoptosis in LTED or exhaustibly anti-estrogens treated breast cancer cells (Jordan *et al*., 2005; Song *et al*., 2001), bone-derived cells (Saintier *et al*., 2006), prostate cancer cells (Robertson *et al*., 1996), neuronal cells (Nilsen *et al*., 2000) and ER-transfected cells (Jiang and Jordan, 1992; Levenson and Jordan, 1994). LTED MCF-7 cells are very sensitive to E2 as such they are stimulated by E2 concentration thousand times lower than the normal MCF-7 cells. This high sensitivity was shown to be the result of upregulation of ERα and MAPK, PI3K, and mammalian target of rapamycin (mTOR) growth factor pathways (Santen *et al*., 2008).

Though the mechanism of tumour regression was unknown previously, recent studies suggest that E2 induces apoptosis in LTED cells through the activation of the Fas death receptor/Fas ligand (FasL) complex (Song *et al*., 2001), as well as alterations in Bcl-2 and the release of cytochrome-c from the mitochondria (Lewis *et al*., 2005; Song *et al*., 2005). But our study has gone further to show for the first time that even high E2 concentration can induce apoptosis in MCF-7 cells as shown in Figure 4B and 4C. The observed data showed that the most potent estrogen in female, E2 causes apoptosis at high concentration even in the MCF-7 cells which is completely a new property of this naturally occurring hormone in vertebrates.

### Variable expression of ERs

Real-time PCR was carried out with the cell line (passage number of around 500) to determine the estrogen receptors ERα, ERβ1 and ERβ2 expression (Figure 5). Gene expression in the samples treated with 0.001µM E2 were compared with the relative gene expression in the untreated control sample to evaluate the effect of increased E2 concentration on ER gene expression. It was observed that ERα and ERβ2 expression increased, whereas the ERβ1 expression was downregulated compared to the control sample (Figure 5). ERα expression was the highest among all the three receptors which showed its importance in the signal transductions involving increased cell proliferation marked by the 0.001µM E2 treatment. A study by Green *et al*. (2009) also showed similar results to the above three ER expression in lobular carcinoma *in situ* (LCIS) of the breast. The steroid hormone E2 with its central role in the regulation of cell differentiation, growth and functioning in the mammary gland, has also a very crucial role in breast cancer progression, invasion and metastasis. It was shown previously (Figure 2 and 3) that 0.001µM E2 treatment increased cell proliferation compared to the untreated control cells and with our current knowledge on the role of ERα in MCF-7 cell proliferation, the observations were consistent with the findings in previous studies (Green *et al*., 2009).

**Figure 5:**
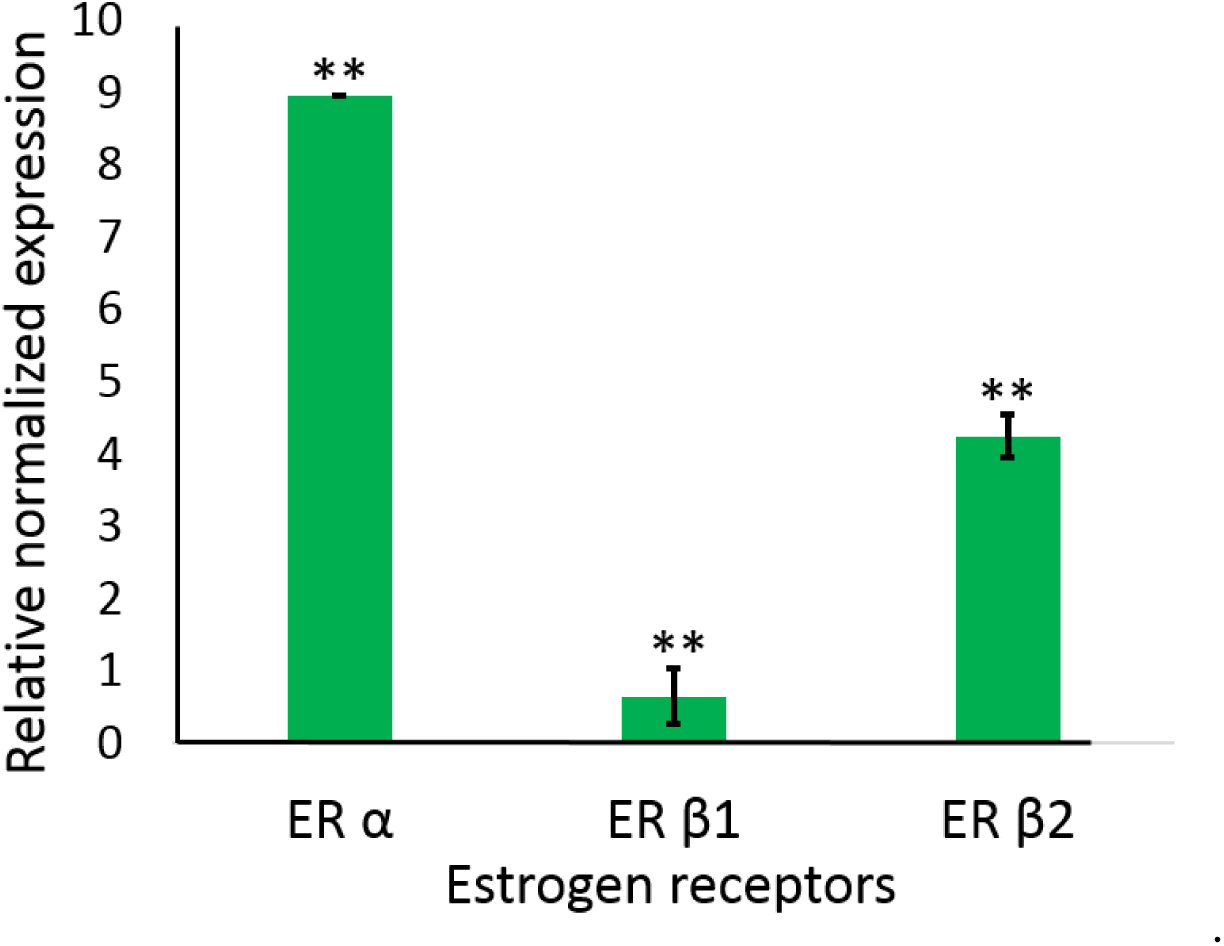
Gene expression of estrogen receptors ERα, ERβ1 and ERβ2 after E2 exposure. Relative gene expression of estrogen receptors ERα, ERβ1 and ERβ2 in response to 0.001µM E2 compared to unstimulated control sample (control=1; Mean ±SEM; n=3; \*\**p*<0.01).

ERβ1 gene was found to be downregulated in the E2 treated cells to the normal untreated cells which supports the findings in the previous studies showing decreased ERβ expression in neoplastic breast cells (Lazennec *et al.,* 2001 and Roger *et al*., 2001). Although it is supported that ERβ is expressed throughout the pathogenesis of breast cancer, the actual role is still controversial. On the contrary, ERβ2 was found to be significantly upregulated like ERα compared to the untreated control samples. This is consistent with the findings showing upregulation of ERβ2 in invasive breast cancer cells compared to the normal cells, with dominant inhibition on the transactivation of ERα dependent genes therefore slowing its mitogenic effects. Furthermore, another study by Kietz *et al*. (2004) has shown a relationship between upregulation of ERα and ERβ2 and which supports our observations.

### Effect of E2 on the expression of ERs

The expression of the three ERs was then evaluated in the new MCF-7 cell line of passage number 149 at two E2 concentration of 0.001µM and 10µM (Fig. 6). Only ERα and ERβ2 were expressed compared to the expression of all three ERs being determined in the old cell line. It was observed that ERs expression increased with the increase in E2 concentration for both ERα and ERβ2. ERα expression was significantly higher in both the E2 concentrations (^*^p < 0.05 vs. control sample; Fig. 6) compared to the control, with the expression at 10µM E2 concentration found to be significantly higher (^+^p < 0.05 vs. 1nM sample; Fig. 6) to that at 0.001µM. Similarly, the ERβ2 expression increased with the increase in E2 concentration, but the expression at 1nM was almost same in contrast to the upregulation of expression at 10µM compared to the control. Moreover, the expression at 10µM was significantly higher (^#^*p* < 0.05 vs. 0.001µM sample; Fig. 6) to that of 0.001µM E2 treated sample. This increase in both ERα and ERβ2 expression with increasing E2 concentration is concordant with the previous finding as discussed previously. Similar results were reported by Li *et al*. (2015), where E1, E2, E3 and E4 were found to significantly upregulate the ERα expression and downregulate the ERβ expression (not expressed in the new cell line; Fig. 6) in a different breast cancer cell line (ZR 75-1) at 1nM concentration compared to the control (Li *et al*., 2015).

**Figure 6:**
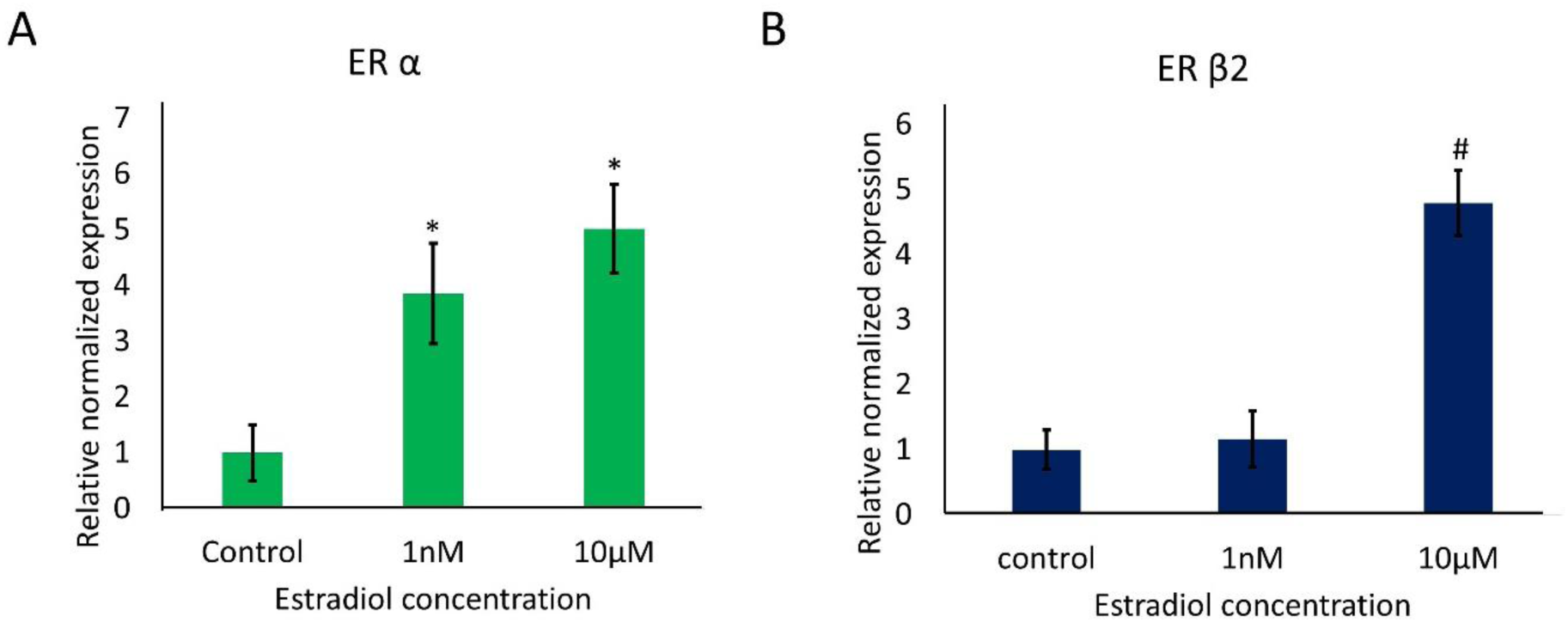
Expression of estrogen receptors, ERα and ERβ2, at two different E2 concentrations 1nM and 10µM. (control=1; Mean ± SEM; n=3, ^*#+^*p*<0.05).

### Effect of subculturing on ERα expression

The ERs expression were considered at different passage number and incubation time with two E2 concentrations. ERβ1 and ERβ2 expression were undetermined in the cells from the sample that was subcultured and incubated with two different E2 concentrations for 16 days (Fig. 7). Interestingly, the only ER subtype expression being determined, ERα, was significantly downregulated (^#^*p* < 0.05 vs. control sample at passage 3=1; Fig. 7) at both the E2 concentrations 1nM and 10µM compared to the control. Similarly, ERα expression was significantly downregulated (^*^*p* < 0.05 vs. 1nM sample at passage 3; Fig. 7) with 1nM E2 at passage 3 compared to passage 0. These results suggest that with increased sub culturing and E2 exposure for longer duration, MCF-7 cells adopt to increased E2 environment by downregulating the ERα expression. This regulation is a classic example of negative feedback in the cells having enough E2 signaling thus trying to balance the intracellular environment by decreasing the amount of ERα present.

**Figure 7:**
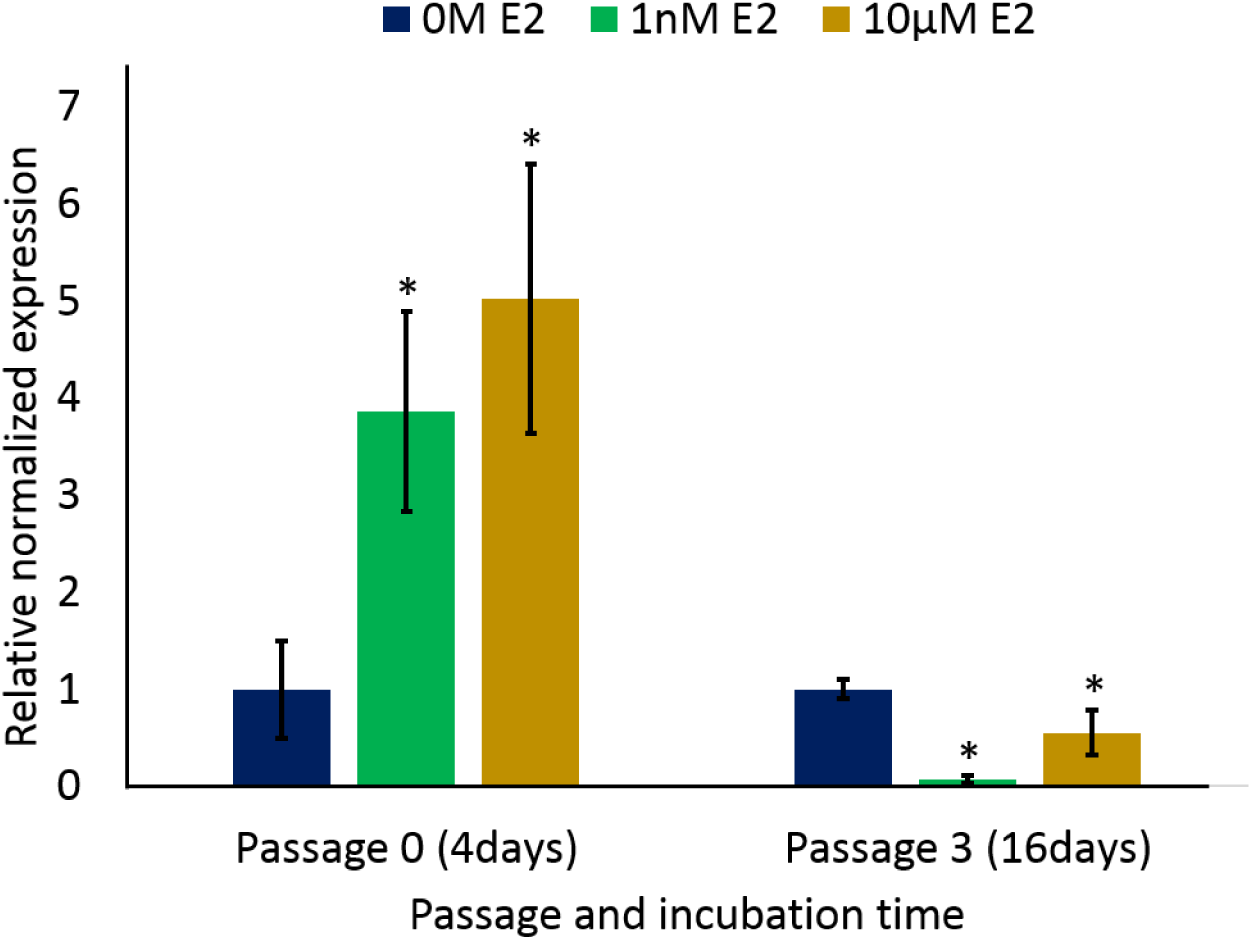
Analysis of ERα gene expression after varied E2 concentration and passage number. Data showing relative ERα gene expression at two different E2 concentrations of 1nM and 10µM at different passage number and incubation time compared to control (control=1; Mean ± SEM; n=2, ^*#+^*p*<0.05).

## CONCLUSION AND FUTURE DIRECTION

It has been previously demonstrated that a very crucial role in breast cancer development, prognosis and occurrence is played by the ER. In addition to the proliferative potential of E2, it was also shown previously to induce apoptosis in LTED or exhaustively anti-estrogens treated breast cancer cells. In conclusion, this study has shown that high concentration of E2 can induce apoptosis in the breast cancer cell line MCF-7, which is a new property of E2 to be uncovered showing its potential application as therapeutics in the treatment of breast carcinoma. This should be an important issue for further research evaluating the clinical benefit of high concentration of E2, as it remains unclear whether this effect can be manifested *in vivo*. Moreover, ERα expression was significantly increased with the increase in E2 concentration, whereas ERβ expression was unchanged at low E2 concentration but upregulated at high E2 concentration compared to the untreated control. MCF-7 cells at passage 3 with continuous E2 exposure for 16 days had significantly downregulated ERα expression compared to both control and cells at passage 0. This study was limited to investigate the effect of both the long term E2 exposure and sub culturing on the receptor expression rather than the effect of sub culturing alone. However, future studies should try to evaluate the effect at high passage number with or without E2 exposure in the breast cancer cells.

## REFERENCES

Altiok, N., Koyuturk, M. and Altiok, S. (2007) JNK pathway regulates estradiol-induced apoptosis in hormone-dependent human breast cancer cells. Breast Cancer Res Treat. Vol.105, pp. 247–254.

Beatson, G.T. (1896) On the treatment of inoperable cases of carcinoma of the mamma: suggestions for a new method of treatment, with illustrative cases. Lancet. Vol. 148:3803, pp. 162–165.

Briske-Anderson, M.J., Finley, J.W. and Newman, S.M. (1997) Influence of culture time and passage number on morphological and physiological development of Caco-2 cells. Proceedings of the Society for Experimental Biology and Medicine. Vol. 214, pp. 248–257.

Chang-Liu, C.M. and Woloschak, G.E. (1997) Effect of passage number on cellular response to DNA-damaging agents: cell survival and gene expression. Cancer Letters. Vol. 26, pp. 77–86.

Esquenet, M., Swinnen, J.V., Heyns, W. and Verhoeven, G. (1997) LNCaP prostatic adenocarcinoma cells derived from high and low passage numbers display divergent responses not only to androgens but also to retinoids. Journal of Steroid Biochemistry and Molecular Biology. Vol.62, pp. 391–399.

Green, A.R., Young, P., Krivinskas, S., Rakha, E.A., Paish, E.C., Powe, D.G. and Ellis, I.O. (2009) The expression of ERa, ERb and PR in lobular carcinoma in situ of the breast determined using laser microdissection and real-time PCR. Histopathology. Vol.54, pp. 419–427.

Gustafsson, J.A. (2003) What pharmacologists can learn from recent advances in estrogen signalling. Trends Pharmacol Sci. Vol.24, pp. 479–485.

Heldring, N., Pike, A., Andersson, S., Matthews, J., Cheng, G., Hartman, J., Tujague, M., Ström, A., Treuter, E., Warner, M. and Gustafsson, J.A. (2007) Estrogen Receptors: How Do They Signal and What Are Their Targets. Physiol Rev. Vol.87, pp. 906–931.

Iwase, H., Zhang, Z., Omoto, Y., Sugiura, H., Yamashita, H., Toyama, T., Iwata, H. and Kobayashi, S. (2003) Clinical significance of the expression of estrogen receptors alpha and beta for endocrine therapy of breast cancer. Cancer Chemother. Pharmacol.Vol.52, pp. S34–S38.

Jiang, S.Y. and Jordan, V.C. (1992) Growth regulation of estrogen receptor negative breast cancer cells transfected with complementary DNAs for estrogen receptor. J Natl Cancer Inst. Vol.84, pp. 580–591.

Jordan, V.C., Lewis, J.S., Osipo, C. and Cheng, D. (2005) The apoptotic action of estrogen following exhaustive antihormonal therapy: a new clinical treatment strategy. Breast. Vol.14, pp. 624–630.

Kerr, J.F.R., Wyllie, A.H. and Currie, A.R. (1972) Apoptosis: A Basic Biological Phenomenon with Wide-ranging Implications in Tissue Kinetics. Br J Cancer. Vol. 26, pp. 239–257.

Kietz, S., Thomsen, J.S., Matthews, J., Pettersson, K., Strom, A. and Gustafsson, J.A. (2004) The Ah receptor inhibits estrogen-induced estrogen receptor beta in breast cancer cells. Biochem. Biophys. Res. Commun. Vol. 320, pp. 76–82.

Kuiper, G.G., Enmark, E., Pelto-Hukko, M., Nilsson, S. and Gustafsson, J-A. (1996) Cloning of a novel estrogen receptor expressed in rat prostate and ovary. PNAS. Vol.93, pp. 5925–5930.

Lazennec, G., Bresson, D., Lucas, A., Chauveau, C. and Vignon, F. (2001) ER beta inhibits proliferation and invasion of breast cancer cells. Endocrinology. Vol.142, pp. 4120–4130.

Levenson, A.S. and Jordan, V.C. (1994) Transfection of human estrogen receptor (ER) cDNA into ER-negative mammalian cell lines. J Steroid Biochem Mol Biol. Vol.51, pp. 229–239.

Lewis, J.S., Meeke, K., Osipo, C., Ross, E.A., Kidawi, N., Li, T., Bell, E., Chandel, N.S. and Jordan, V.C. (2005) Intrinsic mechanism of estradiolinduced apoptosis in breast cancer cells resistant to estrogen deprivation. J Natl Cancer Inst. Vol.97, pp. 1746–1759.

Leygue, E., Dotzlaw, H., Watson, P.H. and Murphy, L.C. (1999) Expression of estrogen receptor β1, β2, and β5 messenger RNAs in human breast tissue. Cancer Research.Vol.59, pp. 1175–1179.

Li, Lv., Wang, Q., Lv, X., Sha, L., Qin, H., Wang, L. and Li, L. (2015) Expression and Localization of Estrogen Receptor in Human Breast Cancer and Its Clinical Significance. Cell Biochem Biophys. Vol.71, pp. 63–68.

Lund, M.J., Butler, E.N., Hair, B.Y., Ward, K.C., Andrews, J.H., Oprea-llies, G., Bayakly, R., O’Regan, R.M., Vertino, P.M. and Eley, J.W. (2010) Age/Race Differences in HER2 Testing and in Incidence Rates for Breast Cancer Triple Subtypes. Cancer. [Online] Available: www.interscience.wiley.com.

Ma, L., Liu, Y., Geng, C., Qi, X. and Jiang, J. (2013) Estrogen receptor β inhibits estradiol-induced proliferation and migration of MCF-7 cells through regulation of mitofusin 2. International journal of oncology. Vol. 42, pp. 1993–2000.

Matthews, J., Wihlen, B., Tujague, M., Wan, J., Strom, A. and Gustafsson, J.A. (2006) Estrogen receptor (ER) beta modulates ER-alpha-mediated transcriptional activation by altering the recruitment of c-Fos and c-Jun to estrogen-responsive promoters. Mol Endocrinol. Vol.20, pp. 534–543.

Matthews, J. and Gustafsson, J.A. (2003) Estrogen signaling: a subtle balance between ER alpha and ER beta. Mol Intervent. Vol.3, pp. 281–292.

Nilsen, J., Mor, G. and Naftolin, F. (2000) Estrogen-regulated developmental neuronal apoptosis is determined by estrogen receptor subtype and the Fas/Fas ligand system. J Neurobiol. Vol.43, pp. 64–78.

Omoto, Y., Kobayashi, S., Inoue, S., Ogawa, S., Toyama, T., Yamashita, H., Muramatsu, M., Gustafsson, J.A. and Iwase, H. (2002) Evaluation of oestrogen receptor β wild-type and variant protein expression, and relationship with clinicopathological factors in breast cancers. European Journal of Cancer. Vol. 38, pp. 380–386.

Pattarozzi, A., Gatti, M., Barbieri, F., Wurth, R., Porcile, C., Lunardi, G., Ratto, A., Favoni, R., Bajetto, A., Ferrari, A. and Florio, T. (2008) 17β-Estradiol Promotes Breast Cancer Cell Proliferation-Inducing Stromal Cell-Derived Factor-1-Mediated Epidermal Growth Factor Receptor Transactivation: Reversal by Gefitinib Pretreatment. Mol Pharmacol. Vol.73, pp. 191–202.

Robertson, C.N., Roberson, K.M., Padilla, G.M., O’Brien, E.T., Cook, J.M. and Kim, C.S. (1996) Fine RL: Induction of apoptosis by diethylstilbestrol in hormone-insensitive prostate cancer cells. J Natl Cancer Inst. Vol.88, pp.908–917.

Roger, P., Sahla, M.E., Makela, S., Gustafsson, J.A., Baldet, P. and Rochefort, H. (2001) Decreased expression of estrogen receptor beta protein in proliferative preinvasive mammary tumors. Cancer Res. Vol. 61, pp. 2537–2541.

Santen, R.J., Song, R.X., Masamura, S., Yue., W, Fan, P., Sogon, T., Hayashi, S., Nakachi, K. and Eguchi, H. (2008) Adaptation to estradiol deprivation causes up-regulation of growth factor pathways and hypersensitivity to estradiol in breast cancer cells. Adv Exp Med Biol. Vol.630, pp.19–34.

Song, R.X., Zhang, Z. and Santen, R.J. (2005) Estrogen rapid action via protein complex formation involving ERalpha and Src. Trends Endocrinol Metab.Vol.16, pp. 347–353.

Song, R.X., Zhang, Z., Mor, G. and Santen, R.J. (2005) Down-regulation of Bcl-2 enhances estrogen apoptotic action in long-term estradioldepleted ER(+) breast cancer cells. Apoptosis. Vol.10, pp. 667–678.

Song, R.X., Mor, G., Naftolin, F., McPherson, R.A., Song, J., Zhang, Z., Yue, W., Wang, J. and Santen, R.J. (2001) Effect of long-term estrogen deprivation on apoptotic responses of breast cancer cells to 17beta estradiol. J Natl Cancer Inst. Vol.93, pp. 1714–1723.

Stewart, B.W. and Wild, C.P. (2014). World Cancer report. [Online] Available: http://www.iarc.fr/en/publications/books/wcr/ [Accessed: 23rd June 2015].

Tremblay, G.B., Tremblay, A., Copeland, N.G., Gilbert, D.J., Jenkins, N.A., Labrie, F. and Giguère, V. (1997) Cloning, chromosomal localization, and functional analysis of the murine estrogen receptor beta. Mol. Endocrinol. Vol.11, pp. 353–365.

